# Combined use of design of experiments and metabolic engineering for optimization of nogalamycin production

**DOI:** 10.1101/804179

**Authors:** Kayla Maki, Marlon Coe, Katelyn Brown, Jennifer Tran, Minji Sohn, S. Eric Nybo

## Abstract

Nogalamycin is an anthracycline antibiotic produced from *Streptomyces nogalater* that exerts its mechanism of action via inhibition of topoisomerase I. Nogalamycin has a unique tetracyclic structure composed of a 7-*O*-glycosidically linked L-nogalose sugar and a C2-C5”-linked L-nogalamine residue that forms an epoxyoxacin ring. Nogalamycin was originally developed as an anticancer agent in the 1970s, however, it exhibited dose-limiting cardiotoxicity. Shortly after the discovery of nogalamycin, the semi-synthetic analogue 7-*O*-methylnogarol, or menogaril, was synthesized and exhibited a mild improvement in cardiotoxicity and anticancer efficacy. Menogaril lacks the 7-*O*-L-nogalose moiety and functions as a topoisomerase II inhibitor, like most anthracyclines, however this agent still proved too toxic for clinical use. Our laboratory is developing a production platform for microbial synthesis of novel nogalamycin analogs useful for treatment of human cancers or as antibiotics. Our initial hypothesis is that overexpression of structural genes responsible for synthesis of TDP-deoxysugar and polyketide precursor substrates, respectively, will increase carbon flux towards nogalamycin production. In this study, we have employed metabolic engineering to enhance nogalamycin production in *Streptomyces nogalater*. In this work, we used an optimized soytone glucose production medium to produce nogalamycin. We also overexpressed copies of structural biosynthetic genes to bolster substrate precursor building blocks for nogalamycin production. First, overexpression of the TDP-glucose synthase and TDP-D-glucose-4,6-dehydratase enzymes (*mtmDE*) resulted in a 50% increase in nogalamycin production (160 mg/L) as compared to wildtype *S. nogalater* (100 mg/L). Secondly, overexpression of the minimal polyketide synthase genes (*snoa123*) resulted in a fourfold production increase in nogalamycin (400 mg/L). This production platform will serve as the fundament for production of nogalamycin analogues for future drug development efforts.

## Introduction

Actinomycetes are industrially indispensable microorganisms that biosynthesize chemicals ranging from antibiotics to antitumor agents and secrete hydrolytic enzymes for the degradation of biochemical polymers. Since wildtype strains do not often exhibit the production phenotype at industrial scale, strain improvement is required. The isolation and characterization of industrial producing strains is accompanied by medium optimization, exemplified by design of experiments (DOE) approaches such as one-factor-at-a-time (OFAT) or factorial design, to identify medium components that augment production ^1^. Though fermentation medium optimization can yield production titer improvements of several-fold, rigorous strain improvement protocols are required to enhance production to hundred milligram or gram per liter scales. Actinomycete strain optimization programs have historically focused on random mutagenesis and high-throughput screening efforts, however, advances in actinomycete genome sequencing and development of recombinant genetic tools have enabled the metabolic engineering of actinomycetes for production of chemicals ^2^.

Previously, Ryu et al. demonstrated that metabolic engineering of carbon flux in actinorhodin producer *Streptomyces coelicolor* M510 (SCP1^−^SCP2^−^Δ*redD*) dramatically enhanced production titer of actinorhodin ^3^. The deletion of *zwf2* encoding glucose-6-phosphate dehydrogenase redirected carbon flux away from the pentose phosphate pathway towards glycolysis and increased rates of synthesis for actinorhodin by 2-fold and biomass 1.6-fold, respectively ^3^. Notably, overexpression of *accA2BE,* encoding the subunits of acetyl-CoA carboxylase, under the control of the *ermE\**p up promoter in *S. coelicolor* M600 (SCP1^−^SCP2^−^) increased actinorhodin production 5-fold from 150 mg/L to 750 mg/L. Recently, Zabala et al. conducted a precursor substrate engineering study in a variety of actinomycetes to enhance production of several different polyketides, including actinorhodin, nogalamycin, chromomycin A3, 8-demethyl-tetracenomycin C, tetracenomycin C, elloramycin, and aclacinomycin ^4^. The authors developed overexpression constructs for the oviedomycin acetyl-CoA carboxylase (i.e. OvmGIH) to enhance carbon flux of acetyl-CoA to malonyl-CoA (Figure 1) in the actinomycete producer strains. Notably, overexpression of *ovmGIH* lead to a 943% increase in actinorhodin production, however, the production titer of anthracyclines was enhanced less than 1-fold via this strategy. For these pathways, this result suggests that regulation of malonyl-CoA metabolism and polyketide biosynthesis might be independently controlled. This observation motivated the present work to extend metabolic engineering studies to include overexpression of polyketide synthase and deoxysugar biosynthesis structural genes in the nogalamycin biosynthetic pathway. We hypothesize that metabolic engineering of the mithramycin *mtmDE* genes encoding TDP-D-glucose and TDP-D-glucose-4,6-dehydratase geneswith the *snoa123* minimal polyketide synthase could more effectively shift carbon flux towards nogalamycin production.

**Figure 1.**
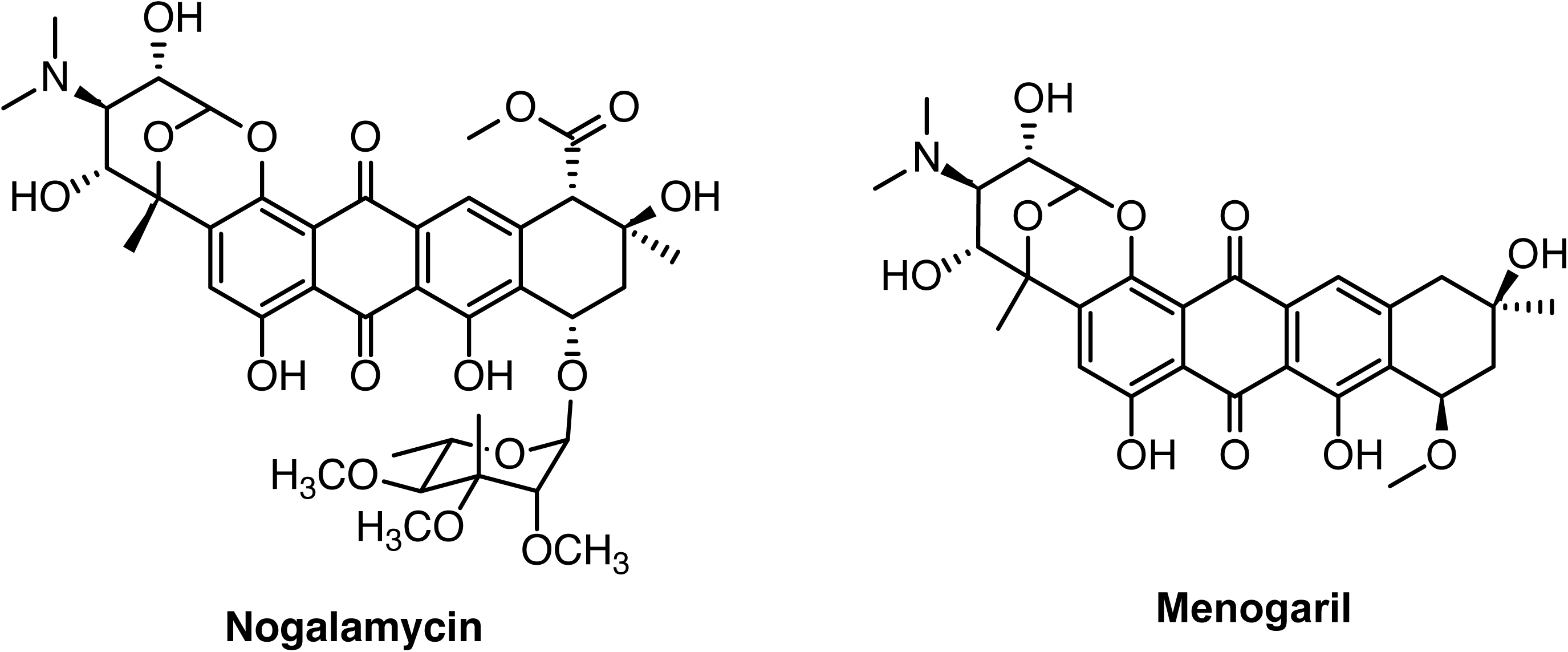
Structures of nogalamycin 299 and menogaril.

Nogalamycin is an anthracycline antibiotic produced by *Streptomyces nogalater* that was discovered in the 1960s ^5^ (Figure 1). Nogalamycin exhibits a unique mechanism of action among anthracyclines in that it inhibits topoisomerase I activity, whereas daunorubicin is primarily a topoisomerase II inhibitor ^6,7^ A less-toxic, semi-synthetic derivative menogaril was developed in the 1970s and briefly enrolled in clinical trials ^8^ (Figure 1). Nogalamycin was recently identified from the National Cancer Institute Compound Library as a lead compound with activity against *Burrelia burgdorferi* Lyme disease and Huntington’s disease ^9,10^. These studies have motivated our present research to investigate new means of production of nogalamycin analogs with improved anticancer efficacy or novel antibiotic properties.

Recently, the biosynthesis of nogalamycin has attracted renewed attention by several groups. The structure of nogalamycin exhibits several interesting structural features, including a heavily oxidized polyketide backbone, an L-nogalose sugar moiety, and an L-rhodosamine sugar with an unusual C5”-C2 carbon-carbon bond ^11^. The nogalamycin polyketide backbone is biosynthesized from 9 malonyl-CoA units and 1 acetyl-CoA unit via the *snoa123* polyketide synthase and undergoes ketoreduction, cyclizations, and oxidation to give rise to nogalamycinone (Figure 2) ^12^. The deoxysugar donor substrates TDP-L-nogalose and TDP-L-rhodosamine are synthesized from 1-phospho-D-glucose via the enzyme SnogJ to form NDP-D-glucose and NDP-4-keto-6-deoxy-D-glucose (Figure 2). Nogalamycinone undergoes subsequent glycosylation, oxidation, and *O*-methylation reactions that and These investigations have elucidated the mechanisms of several polyketide tailoring enzymes, such as the unique snoaL2/snoaW C-1 hydroxylase ^13^, the unique C5”-C2 carbon-carbon bond forming SnoK oxygenase ^14^, and two glycosyltransferases responsible for transfer of TDP-L-nogalose and TDP-L-rhodosamine to the nogalamycinone aglycone ^15^.

**Figure 2.**
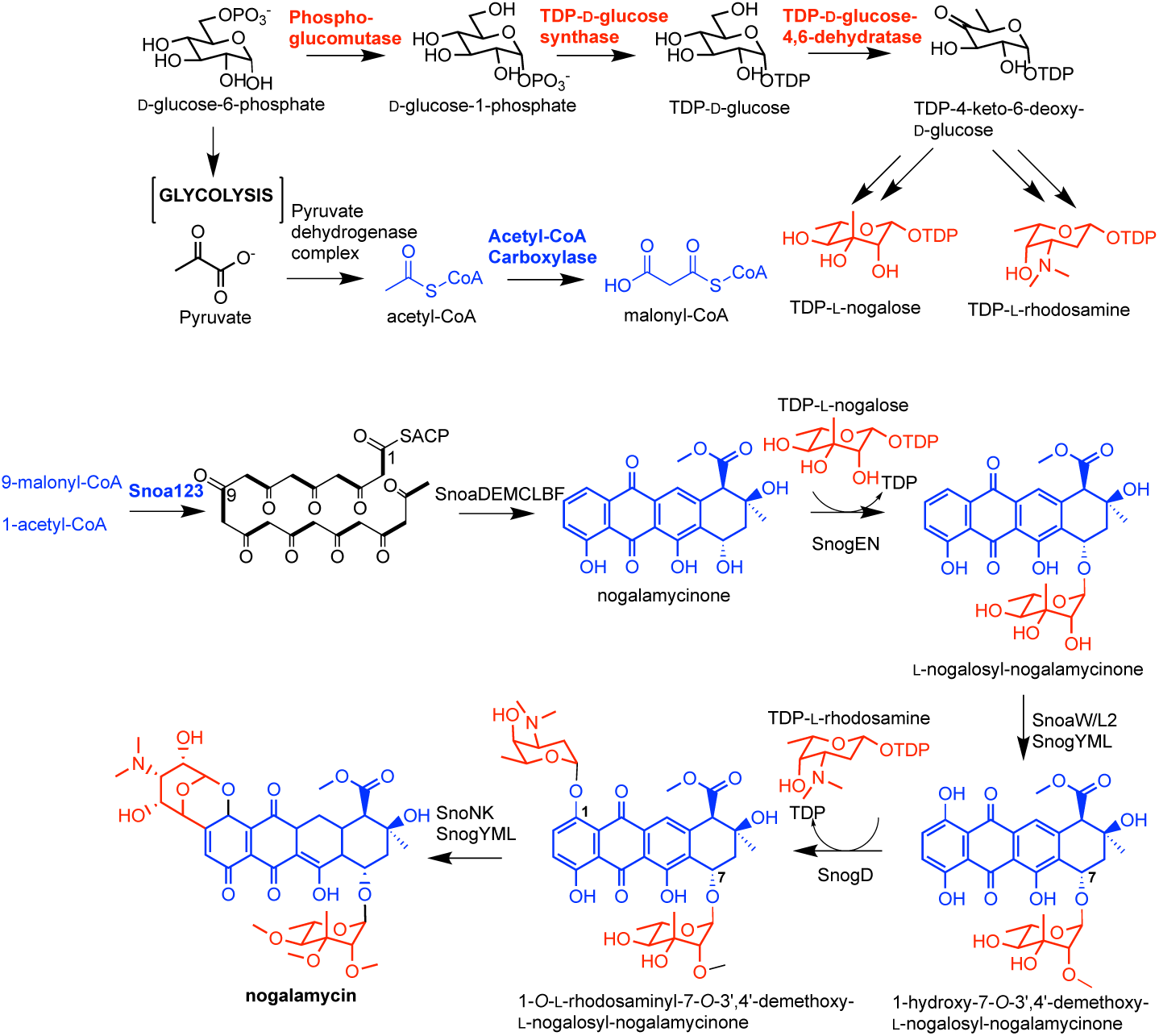
The nogalamycin biosynthetic pathway with enzymatic steps targeted for metabolic engineering highlighted in **red** (polyketide acceptor substrate) or **blue** (TDP-deoxysugar donor substrate). D-glucose-6-phosphate is converted by phosphoglucomutase to D-glucose-1-phospahte, which is converted by the actions of TDP-D-glucose synthase and TDP-D-glucose-4,6-dehydratase to the key intermediate TDP-4-keto-6-deoxy-D-glucose. TDP-4-keto-6-deoxy-D-glucose is converted to TDP-L-nogalose and TDP-L-rhodosamine. The acetyl-CoA carboxylase complex catalyzes carboxylation of acetyl-CoA to form malonyl-CoA, which are both building blocks for the Snoa123 minimal polyketide synthase. The Snoa123+SnoaDEMCLBF polyketide synthase forms nogalamycinone. Additional post-PKS tailoring enzymes, including glycosyltransferases, oxygenases, and *O*-methyltransferases generate nogalamycin.

In this work, we developed a combined design of experiments and metabolic engineering approach to modulate the production of nogalamycin. First, we applied one-factor-at-a-time (OFAT) analysis of GYM, NDYE, and SG liquid media for production of nogalamycin. We supplemented these media with different carbon sources, including glucose, glycerol, starch, and maltose and different nitrogen sources, including tryptone, yeast extract, peptone, and soytone. Secondly, we metabolically overexpressed *mtmDE* and *snoa123* polyketide synthase in *S. nogalater* for enhanced production of nogalamycin. To enhance *in vivo* biosynthesis of nogalamycin, we overexpressed *mtmDE* (i.e. encoding NDP-D-glucose synthase and NDP-D-glucose-4,6-dehydratase) from the mithramycin pathway and the *snoa123* nogalamycin polyketide synthase genes. The genes were expressed under the control of the strong erythromycin resistance promoter *ermE*p* in a high-copy number plasmid pWHM3 and the resulting expression vectors were transformed into *Streptomyces nogalater* along with empty plasmid vector control.

## Methods and Materials

### Bacterial strains and growth conditions

*Streptomyces nogalater* NRRL 3035 was obtained from the USDA NRRL Culture Collection (Peoria, IL). *Streptomyces nogalater* was plated on mannitol soya flour agar for 3 days at 28°C until sporulation as described previously ^16^ Spores were inoculated into 5 mL of tryptic soy broth medium (TSB) and grown for 2 days at 30°C to prepare a seed culture. *S. nogalater* was transformed via intergeneric conjugation and *E. coli* ET12567/pUZ8002 was used as a conjugation donor as previously described ^17^. TSB cultures were inoculated 1% v/v into 3 mL culture tubes or 50 mL shake flask cultures of GYM, NDYE, or SG media as previously reported ^18^.

### Construction of expression plasmids

pWHM3 was used as a host plasmid for the expression vectors described in this work. All of the promoters and genes in this work were cloned as BioBrick fragments as previously described ^19^. The *ermE\**p was synthesized as a double stranded BioBrick gBlock (IDT DNA). *Snoa123* was amplified via polymerase chain reaction from *S. nogalater* NRRL 3035 genomic DNA. *MtmDE* were amplified via PCR from plasmid pFL492 (Jurgen Rohr, University of Kentucky) ^20^. PCR products were cloned using the Zero Blunt™ TOPO^®^ cloning kit following the manufacturer’s instructions (Invitrogen). All molecular biology reagents were purchased from New England Biolabs. Plasmids were purified using the Wizard Miniprep SV kit.

### Analysis of cell biomass and nogalamycin production

*S. nogalater* cultures were sampled every 24 hours to determine biomass and nogalamycin production titer. One milliliter samples were collected in a microcentrifuge tube, centrifuged to collect the cell pellet, then the cell pellet was dried. Biomass was calculated by weighing the vessel and determining grams of dry cell weight (g DCW). One milliliter of culture extract was purified via C18 solid phase extraction cartridge (50 mg HyperSep C18, ThermoFisher) at 5 millibar of pressure. The retained material was washed with water and eluted in 1 mL of methanol. HPLC-MS analysis of nogalamycin was carried out at the Ferris State University, Shimadzu Core Laboratory on a Shimadzu LCMS-8040 Liquid Chromatography Platform with Triple Quad Mass Spectrometer Interface. UV-vis detection of nogalamycin was carried out at 254 and 478 nm wavelength and nogalamycin parent mass fragment [M+H]^+^ 788 *amu.* Extracts were chromatographed on a reversed phase column (Prodigy™ 5 µm ODS-3V C_18_, 4.6 mm x 250 mm), with isocratic gradient of 40:60 acetonitrile and water containing 0.1% formic acid for 20 minutes. A standard curve of nogalamycin from 0-100 μg/mL was quantified via HPLC-MS to obtain linear plots.

## Results

### One-factor-at-a-time Medium Optimization for Production of Nogalamycin

To determine biomass production and nogalamycin titer of *S. nogalater* NRRL 3035 in a variety of production media, the wildtype strain was first grown in triplicate 50 mL GYM, NDYE, and SG liquid media shake flask fermentations for 4 days (Figure 2). *S. nogalater* exhibited the highest biomass production and nogalamycin titer in SG liquid media (120 mg/L and 17 gCDW/L), followed by NDYE media (80 mg/L and 15 gCDW/L), and last GYM media (20 mg/L and 12 gCDW/L) (Figure 3). This experiment established a baseline nogalamycin production titer in each production medium. Next, we supplemented each production medium with different carbon sources: 2% w/v glucose, 2% v/v glycerol, 2% w/v maltose, and 2% w/v starch and different nitrogen sources: 2% w/v soytone, 2% w/v tryptone, 2% w/v peptone, 2% w/v yeast extract. The wildtype *S. nogalater* was grown in double triplicate 3 mL culture tube fermentations of each type of medium with and without carbon source or nitrogen source supplementation for 3 days (Figure 3). We compared the production of nogalamycin between supplemented media and basal media and performed untailed T-test statistical analysis to determine statistical significance.

**Figure 3.**
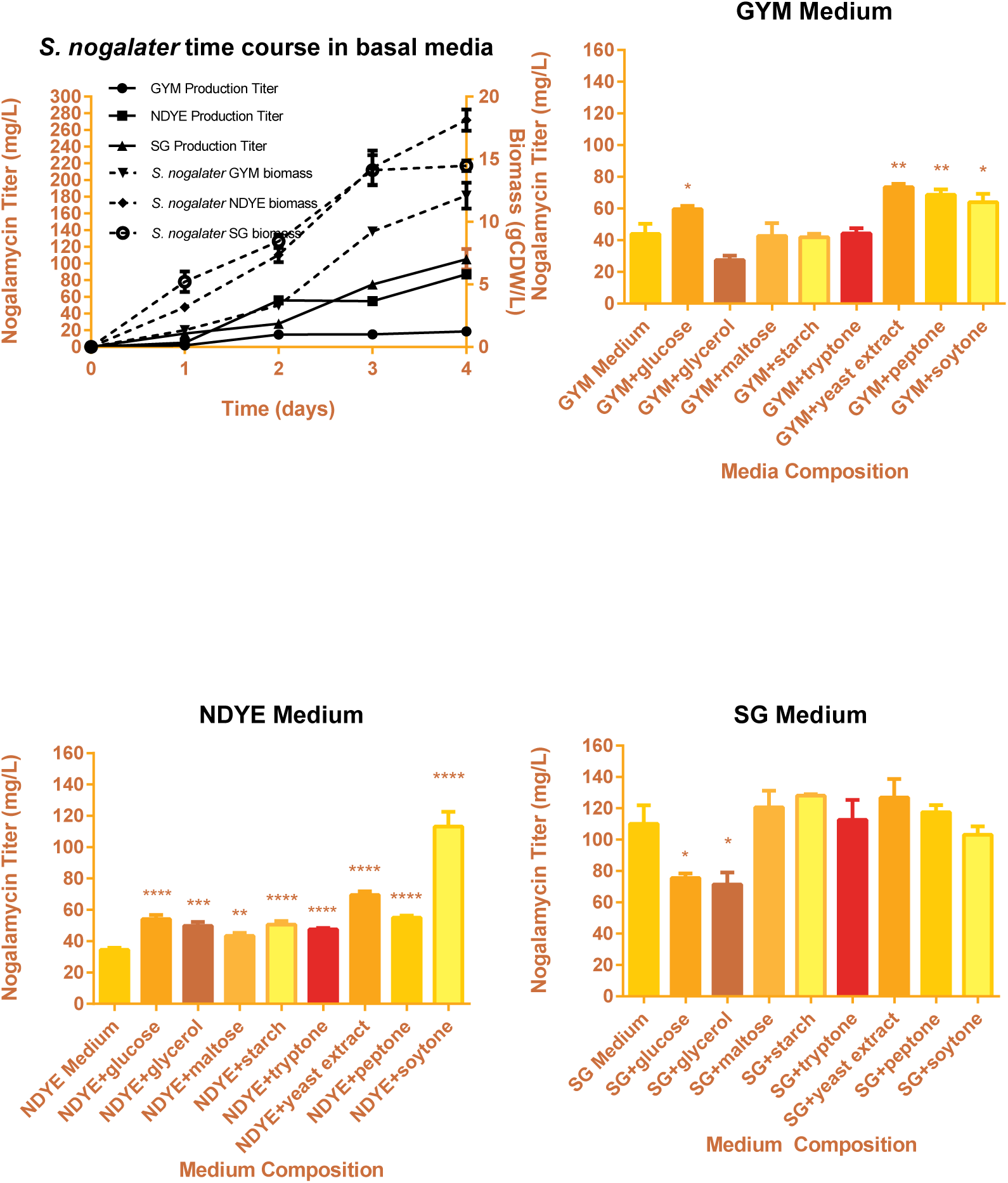
Medium optimization for production of nogalamycin. *Streptomyces nogalater* was grown in GYM, NDYE, and SG liquid media in a time course (upper left panel). *Streptomyces nogalater* was grown in GYM, NDYE, and SG liquid medium supplemented with 2% glucose, glycerol, maltose, tryptone, yeast extract, peptone, or soytone. Medium with supplementation was compared to the base medium via two-sided T-test. *indicates statistically significant comparison *p*<0.05, ** indicates *p*<0.01 and is very significant, *** indicates *p*<0.001, and **** indicates *p*<0.0001.

In general, the culture tube fermentations exhibited lower production titers overall, most likely due to poorer oxygenation. Supplementation of GYM media with glucose, yeast extract, peptone, or soytone resulted in statistically significant increase of nogalamycin production by 33% as compared to GYM media (60 mg/L, Figure 2). Supplementation of NDYE with all carbon and nitrogen sources tested resulted in statistically significant increase in production of nogalamycin of 1 to 4-fold (60 mg/L to 120 mg/L). Supplementation of SG media with glucose and glycerol lead to a statistically significant 20% decrease in nogalamycin production titer. Supplementation with other carbon and nitrogen sources increased production slightly, though the result was not statistically significant.

### Metabolic engineering of nogalamycin production

SG media was selected as an optimal fermentation medium for production of nogalamycin. We next transformed *S. nogalater* with several overexpression constructs to determine potential rate-limiting steps of nogalamycin metabolism (Figure 4). First, we introduced constructs overexpressing the *mtmDE* genes to enhance carbon flux to NDP-D-glucose and NDP-4-keto-6-deoxy-D-glucose. *S. nogalater* (pWHM3-*mtmDE*) produced a maximum titer of 160 mg/L nogalamycin, which was a 50% increase to the production titer of the control strain *S. nogalater* (pWHM3) (100 mg/L) (Figure 5). This indicates that synthesis of NDP-deoxysugar substrates limits nogalamycin production. More importantly, overexpression of the *snoa123* polyketide synthase resulted in a fourfold increase in nogalamycin production (400 mg/L) (Figure 5). This result demonstrated that synthesis of the polyketide core from malonyl-CoA limits overall carbon flux through the pathway.

**Figure 4.**
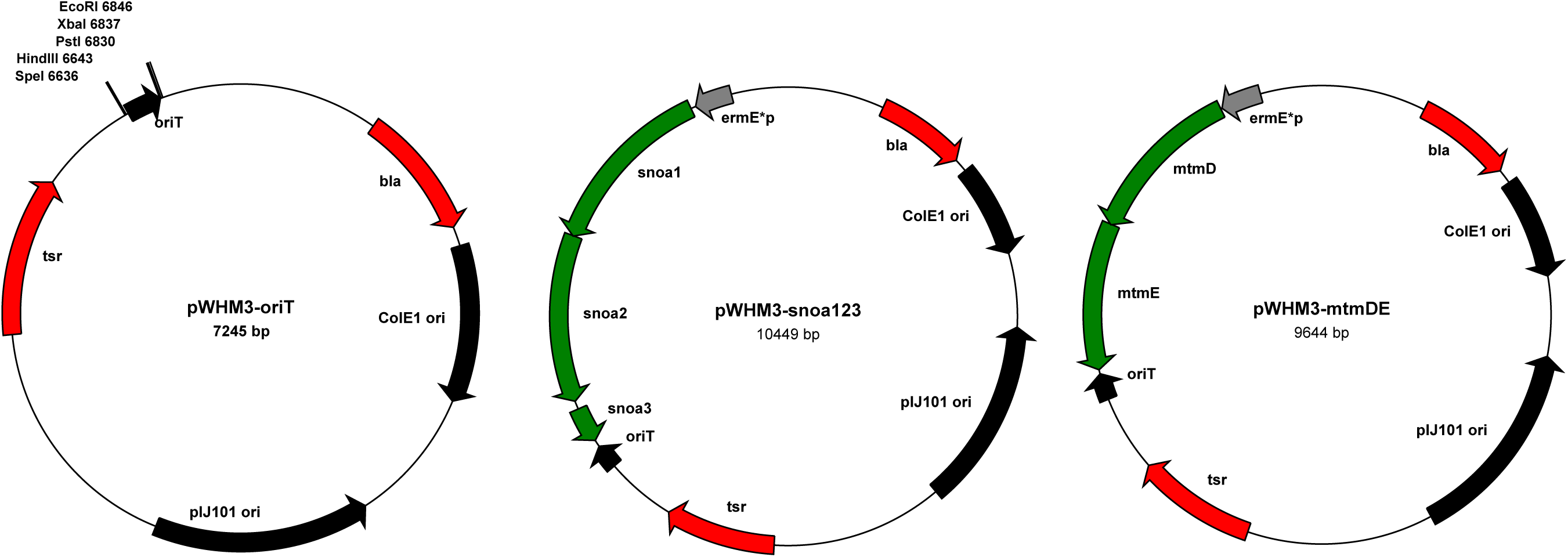
Plasmid vectors for metabolic engineering of nogalamycin production. pWHM3-*oriT* is a high-copy number *E. coli-Streptomyces* shuttle vector for overexpression of biosynthetic genes from the constitutive *ermE*p* promoter. Constructs include pWHM3, pWHM3-*mtmDE* (e.g. TDP-D-glucose synthase and TDP-D-glucose-4,6-dehydratase from *S. argillaceus*), and pWHM3-*snoa123* from (e.g. minimal polyketide synthase from *S. nogalater*).

**Figure 5.**
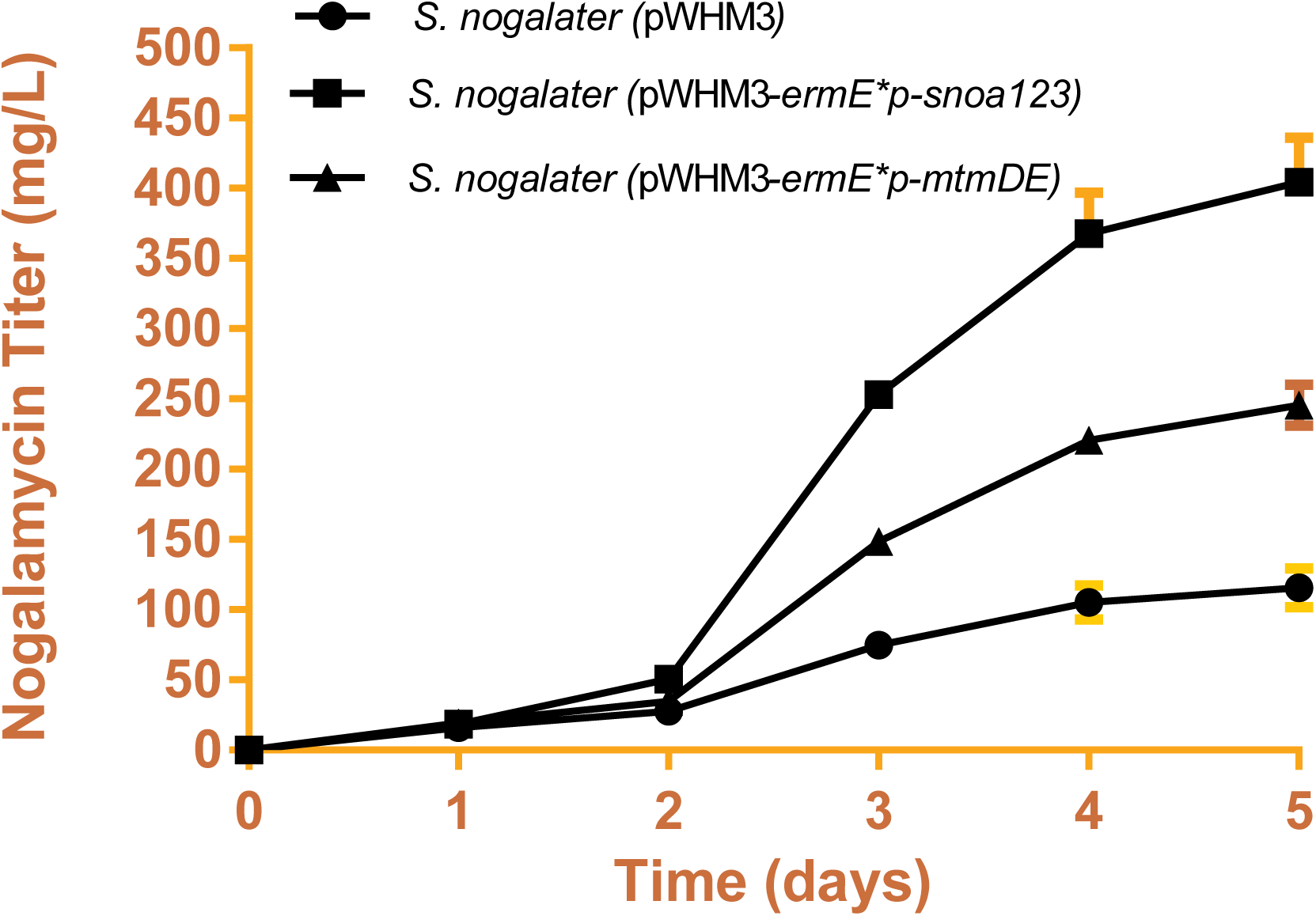
Time course experiment of metabolically engineered *S. nogalater* lines. *Streptomyces nogalater* was transformed with pWHM3-*oriT* (closed circle), pWHM3-*ermE*p-snoa123* (closed square), or pWHM3-*ermE*p-mtmDE* (closed triangle) and grown for five days in liquid SG medium. Nogalamycin production was determined daily (mg/L). Error bars reflect S.E.M. of triplicate biological experiments.

## Discussion

OFAT design-of-experiments and metabolic engineering are synergistic approaches to optimize nutrient availability and genetic regulation of metabolic pathways. In this work, we grew *S. nogalater* in three laboratory production media and improved nogalamycin production threefold (120 mg/L) (Figure 3). Supplementation of the growth media resulted in contextual improvements in GM and NDYE media experiments. Glucose, glycerol, maltose, and starch carbon sources all improved nogalamycin production when supplemented into NDYE. Similarly, tryptone, yeast extract, peptone, and especially soytone improved nogalamycin production when supplemented into NDYE. NDYE is a defined medium that was originally developed for production of doxorubicin in *Streptomyces peucetius*, a pathway that is inhibited by carbon catabolite repression ^21^. This experiment demonstrates that NDYE is not optimized for nogalamycin production in *Streptomyces nogalater*, possibly due to the fact that it lacks a readily accessible carbon or nitrogen source, and probably because much of the carbon and nitrogen are directed towards biomass synthesis (Figure 2). This result also demonstrates that *Streptomyces nogalater* is capable of using additional carbon and nitrogen sources for polyketide biosynthesis. On the other hand, GYM media experiments demonstrated a statistically significant increase in nogalamycin production when they were supplemented with additional glucose, yeast extract, peptone, or soytone. These experiments demonstrate that *S. nogalater* benefits from additional supplementation with glucose, its preferred carbon source, and additional nitrogen sources. Nutrient supplementation into SG liquid medium failed to increase production of nogalamycin, and in two cases, addition of glucose and glycerol decreased production. One possible explanation for this could be that additional glucose supplementation is directed towards biomass production and not nogalamycin production, which is reflected in the high biomass production of *S. nogalater* in SG liquid medium (17 gCDW/L).

Most importantly, substrate precursor metabolic engineering was demonstrated to further take advantage of additional nutrient supplementation in SG liquid medium. When *Streptomyces nogalater* was transformed with empty plasmid vector, pWHM3, no additional increase in nogalamycin production was observed. Engineering of *mtmDE* resulted in a 50% increase in nogalamycin production to 160 mg/L when compared to the control strain, most likely due to the increase of TDP-deoxysugar precursors in the synthesis of TDP-_L_-rhodosamine and TDP-_L_-nogalose (Figure 5). Overexpression of the *snoa123* minimal polyketide synthase genes lead to a fourfold increase of nogalamycin production (400 mg/L). Ultimately, these experiments demonstrate the synergy of combining traditional one-factor-at-a-time fermentation optimization and substrate precursor engineering for titer improvement in important natural products.

